# A closed genome of *Streptococcus agalactiae* ST347/II reveals a composite integrative element coupling nisin P biosynthesis to *erm(B)* and tet(O) in Dominican group B *Streptococcus*

**DOI:** 10.64898/2026.10.29.762887

**Authors:** Leslys García-Encarnación, Dolinsky S. De La Cruz-Caminero, Hansel B. López-Chevalier

**Affiliations:** Instituto Especializado de Estudios Superiores Loyola (IEESL), San Cristóbal, Dominican Republic; Facultad de Ciencias, Universidad Autónoma de Santo Domingo (UASD), Santo Domingo, Dominican Republic

**Keywords:** *Streptococcus agalactiae*, group B Streptococcus, nisin P, lantibiotic, integrative conjugative element, *erm(B)*, macrolide resistance, Dominican Republic

## Abstract

Clindamycin is the alternative for intrapartum prophylaxis against group B *Streptococcus* in penicillin-allergic women, and whether it works depends on which lineages carry the resistance and on the mobile element that moves it. No Dominican group B *Streptococcus* genome has been closed. A *Streptococcus agalactiae* isolate from residual diagnostic material was sequenced twice on Oxford Nanopore, annotated, and screened against 83 genomes from a published Dominican maternal colonisation survey. The assembly is one circular chromosome of 2,066,859 bp with all seven rRNA operons resolved: ST347/CC19, serotype II, carrying *erm(B), tet(O), mef(J)* and *msr(I)*. Two integrative conjugative elements were delimited: one of ~34 kb related to Tn5252 carrying *erm(B)* and *tet(O)*, another of ~35 kb at a tRNA-Lys gene carrying *mef(J)* and *msr(I)*. An eleven-gene class I lanthipeptide cluster lies 96 bp from the first element, on the same strand and with no annotated feature between them; its precursor encodes a 31-residue core identical to nisin P. The complete operon was found in 12 of the 83 genomes, 13 of 84 (15.5%) with the isolate. *tet(O)* was present in all 12 carriers and none of the 71 others (Fisher p = 1.03 x 10^-14), *erm(B)* in 12 of 12 against 3 of 71 (p = 4.70 x 10^-12). All eleven ST1 genomes lack both the operon and the nisin resistance protein; all 15 ST1010/IV genomes carry *aac(6’)-Ie*/*aph(2’’)-Ia*. Clindamycin resistance is therefore tightly associated with a lantibiotic-encoding element and contiguous with it in the one closed genome; in the drafts the relationship is statistical, and co-occurrence is not linkage. The ST1 absence predicts lineage-wide susceptibility, testable by deferred antagonism.

## Introduction

*Streptococcus agalactiae* remains the leading cause of earlyonset neonatal sepsis and meningitis, and intrapartum antibiotic prophylaxis of colonised mothers is the intervention that prevents it (1, 2, 3). Penicillin is the first-line agent and the species has no acquired beta-lactamase, so prophylaxis fails for a different reason: in women with reported penicillin allergy, clindamycin is the alternative (3, 4), and clindamycin resistance is acquired, mobile and increasing (5, 6). Failure of prophylaxis attributable to clindamycin-resistant GBS has been documented in early-onset disease (7). Whether a given lineage carries that resistance therefore determines whether the alternative regimen works, which makes the genetic basis of macrolide and lincosamide resistance in a local GBS population a question of direct clinical consequence rather than surveillance alone.

Antimicrobial resistance was associated with 4.71 million deaths and directly attributable to 1.14 million in 2021, and tropical Latin America is among the regions where annual attributable deaths rose most between 1990 and 2021 (8); the burden falls disproportionately on low- and middle-income countries (9). Of ten antibacterials approved between 2017 and 2023, only two were judged innovative and none belonged to a new class (10), so interest has returned to bacteriocins and to ribosomally synthesised and post-translationally modified peptides (RiPPs), which act on targets the current classes do not (11). Nisin and its relatives bind lipid II, combining inhibition of cell wall synthesis with pore formation (12).

Among RiPPs, nisin and its natural variants are the best characterised class I lanthipeptides. Variants have been described from *Lactococcus lactis, Streptococcus uberis* (13), *S. hyointestinalis, S. equinus, S. salivarius, Staphylococcus capitis* and gut anaerobes, differing mainly in the hinge region, the flexible segment between rings C and D that governs activity and target spectrum and that protein engineering has repeatedly identified as the decisive determinant (14, 15, 16). Nisin P is the shortest natural variant at 31 residues, with an AIK hinge, and reported activity against methicillinresistant *Staphylococcus aureus*, vancomycin-resistant enterococci and penicillin-resistant *S. pneumoniae*; it was first identified in *S. pasteurianus*, later found in *S. suis*, and demonstrated to be produced by a single human faecal *S. agalactiae* isolate, DPC7040, the only GBS producer reported to date (17). *S. agalactiae* also carries a chromosomal nisin resistance protein, a membrane-associated protease that cleaves the peptide rather than shielding the membrane as producer immunity proteins do (18), so production and resistance can coexist within the species.

Two gaps motivated this work. First, the genetic context of resistance determinants cannot be resolved in fragmented assemblies: integrative conjugative elements span tens of kilobases, their boundaries lie in repeats, and a determinant on a contig edge cannot be assigned to an element. Second, the only genomic description of Dominican GBS is a 2021-2022 survey of rectovaginal colonisation in 350 near-term pregnant women at a tertiary hospital in Santo Domingo, which found 26.6% colonisation and sequenced 93 isolates (19). That study reported serotypes, sequence types and resistance determinants by class. It did not examine biosynthetic gene clusters, did not resolve the genetic context of the resistance determinants it found, and could not do so: every one of its assemblies is a draft. We sequenced one diagnostic isolate to closure, used it to resolve the architecture its draft counterparts cannot, and then used that architecture as a query against the national collection.

## Results

### A closed ST347/II chromosome with four acquired resistance determinants

Both libraries assembled into a single circular contig of exactly 2,066,859 bp. Alignment of the two assemblies gave 100% aligned bases at 99.99% average identity, differing at five positions: three single-nucleotide substitutions and two indels. All five are attributable to basecalling rather than to sequence variation. The two indels lie within homopolymers, a run of twelve thymines and a run of nine adenines, which is the characteristic error mode of nanopore basecalling. At the three substituted positions both read sets carried both bases at normal depth, and the minor-allele fraction differed between positions — 29.4%, 46.6% and 55.8% pooled across libraries, heterogeneous at χ^2^ = 59.25, 2 df, p = 1.4 × 10^−13^ — which excludes the single mixing ratio that two co-resident clones would impose. The five positions are reported as unresolved rather than curated to one state. The full analysis, including the control that excludes partition of a single read set, is given in Materials and Methods, and per-position read counts in Supplementary Table S11.

| **Gene** | **Coordinates** | **Identity** | **Class** |
|---|---|---|---|
| *erm*(B) | 318,281-319,018 | 99.59% | Lincosamide, macrolide, streptogramin |
| *tet*(O) | 328,128-330,047 | 100% | Tetracycline |
| *mef*(J) | 1,137,722-1,138,915 | 100% | Macrolide |
| *msr*(I) | 1,139,029-1,140,486 | 99.79% | Macrolide, streptogramin |

No beta-lactam resistance determinant was found, consistent with the species. Bakta annotates a protein at 282,559-283,710 as a beta-lactamase on the basis of fold homology; AMRFinderPlus did not call it in either run, and it is not reported here as a resistance determinant.

The annotation also places a short open reading frame at 319,143-319,226, immediately downstream of *erm*(B) and on the same strand, encoding the rRNA methylase leader peptide. This is the translational attenuator that makes *erm*(B) expression inducible (21). Its presence predicts an inducible MLSb phenotype, in which clindamycin can appear susceptible by routine disc or broth testing while macrolide exposure derepresses the methylase. The prediction is directly testable by a double-disc diffusion (D) test on this isolate (22), and it carries clinical weight precisely because clindamycin is the agent used for intrapartum prophylaxis in women reporting penicillin allergy.

| **Feature** | **Value** |
|---|---|
| Chromosome length | 2,066,859 bp |
| Contigs | 1, circular |
| GC content | 35.5% |
| Coding density | 90.0% |
| CDS | 1,984 (192 hypothetical) |
| tRNA | 80 |
| tmRNA | 1 |
| rRNA genes | 21, in 7 complete operons |
| ncRNA | 13, plus 18 ncRNA regions |
| CRISPR arrays | 1 |
| oriC/oriV | 2 |
| Assembly gaps | 0 |

The 21 rRNA genes resolve into seven complete operons. Because these operons are near-identical repeats longer than a short-read insert, they are among the regions a shortread assembly cannot resolve, and the 82 contig-level comparison genomes offer no equivalent view of them. Annotation also detected two origins of replication in the single circular chromosome.

The isolate is sequence type 347, clonal complex 19, capsular serotype II. Three other Dominican ST347 draft genomes are deposited in public databases, and all three are serotype II, so ST347/II is a coherent local lineage rather than a single isolate; none of the three is closed.

Four acquired resistance determinants were found, and the same four were returned by three independent calls: AMRFinderPlus 3.12.8 with its 2024 database, AMRFinder-Plus 4.2.7 with database 2026-08-07.1, and the AMRFinder expert system inside Bakta 1.12.2. Coordinates and identities were identical between the two AMRFinderPlus runs, so the resistome call does not depend on database version.

A putative nisin resistance protein, a membrane-associated protease belonging to the species’ chromosomal complement, is annotated at 2,035,261-2,036,223.

### Two integrative conjugative elements, and a lanthipeptide cluster contiguous with one of them

The four resistance genes sit on two separate elements, not one (Figure 1). Integrative conjugative elements excise from the chromosome, transfer by conjugation and integrate into the recipient (23, 24); neither assignment below is possible in a fragmented assembly.

**Figure 1.**
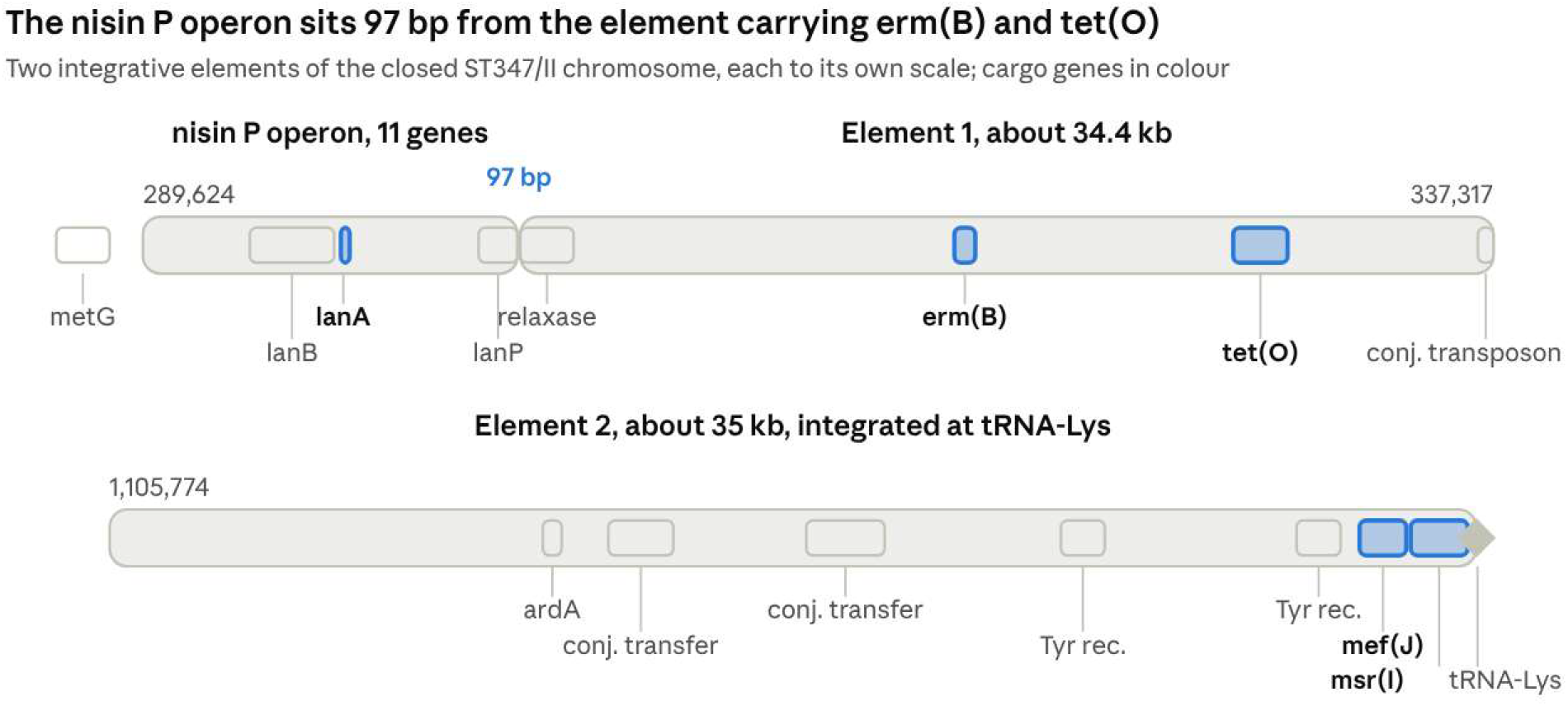
The nisin P operon lies 96 bp from the integrative element carrying *erm(B)* and tet(O). Two integrative conjugative elements of the closed ST347/II chromosome, each drawn to its own scale; cargo resistance genes and the precursor gene are highlighted. Element 1 (approximately 34.4 kb) carries *erm(B)* and tet(O); element 2 (approximately 35 kb) carries mef(J) and msr(I) and is bounded on the right by the tRNA-Lys gene into which it integrated.

#### Element 1, approximately 302,951 to 337,317 (34.4 kb), carrying *erm*(B) and *tet*(O)

The region opens with a relaxase and mobilisation nuclease (302,951-304,834) and a MobC-family mobilisation protein (304,821-305,186), followed by a protein annotated as Tn*5252* (305,189-305,551) (25, 26), the conjugative transposon of *S. pneumoniae*. Downstream lie a 3’-5’ DNA helicase, a *pezA* antitoxin, a CHC2 zinc-finger primase, a protein annotated simply as an integrative and conjugative element protein (314,243-314,542), an SNF2-family helicase and an AAA-ATPase. *erm*(B) sits at 318,281, a transposase at 319,354, a ParB/Spo0J partition protein at 322,296 and a plasmid recombination enzyme at 323,158. *tet*(O) sits at 328,128, followed by *tnpV* at 330,414, two large proteins of 2.0 and 3.9 kb carrying helicase and methyltransferase domains, and a conjugative transposon protein at 336,766.

The element carries *tet*(O) rather than the *tet*(M) of canonical Tn*916*, and the Tn*5252* and TnpV signatures place it in a different lineage of streptococcal conjugative elements. The presence of a relaxase, a mobilisation protein, a partition system, a toxin-antitoxin pair and conjugative transposon proteins indicates a self-transmissible element rather than a passively mobilised one.

#### Element 2, approximately 1,105,774 to 1,140,700 (35 kb), carrying *mef(J)* and *msr(I)*

This element has the more complete and more recognisable conjugation apparatus: three conjugal transfer proteins at 1,118,520, 1,119,771 and 1,123,581, the last 2.0 kb and consistent with an FtsK/SpoIIIE-family translocase; a C51/CHAP-domain peptidase at 1,126,077, the cell-wall hydrolase such elements use to open a conjugation pore; an antirestriction protein ArdA at 1,116,846, the Tn916 orf18 product, which inhibits the recipient’s type I restriction-modification system and so protects incoming DNA (27); a *relK* toxin with its antitoxin at 1,117,954-1,118,467; and Cro/CI-family regulators. Two tyrosine recombinases are present, at 1,130,088 and 1,136,110, indicating an element within an element. *mef(J)* and *msr(I)* lie at 1,137,722 and 1,139,029, at the distal end of the region, and a tRNA-Lys gene begins at 1,140,702. Integration at a tRNA gene is the canonical attachment site for streptococcal integrative conjugative elements of the Tn916/ICESt3 family (28); it fixes the righthand boundary.

A third conjugation module without resistance cargo lies between 706,847 and 722,218, carrying a transposase and an integrase. The genome therefore contains at least three integrative elements of related families, two of which carry acquired resistance. The second of the two origins of replication detected in the chromosome is most plausibly that of one of these elements, several of which can replicate autonomously once excised (20).

#### Mobilome prediction

geNomad reported no plasmids and a single provirus at 1,854,192-1,866,183, spanning 11,992 bp with 14 genes, one integrase, a virus score of 0.8994 and marker enrichment of 10.09, assigned to the *Caudoviricetes*. It detected no viral hallmark genes, that is, no capsid, portal or terminase. At 12 kb with no structural module, this is best described as a prophage remnant or a phage-derived integrative region rather than an inducible prophage; complete GBS prophages are typically 30 to 50 kb. Neither resistance locus lies within or near it, which, together with the absence of plasmids, establishes that both are chromosomal and element-borne.

#### A complete nisin P operon, 96 bp from the resistance element

Immediately adjacent to element 1, at lower coordinates and on the same strand, lies a complete class I lanthipeptide biosynthetic cluster of eleven genes, spanning 289,624 to 302,854. Its last gene, the leader-processing protease, ends at 302,854, and the relaxase that opens the resistance element begins at 302,951, leaving 96 bp between them. Both are annotated on the minus strand, and the annotation places no feature in the intervening sequence, so the two loci are contiguous and co-oriented rather than merely close.

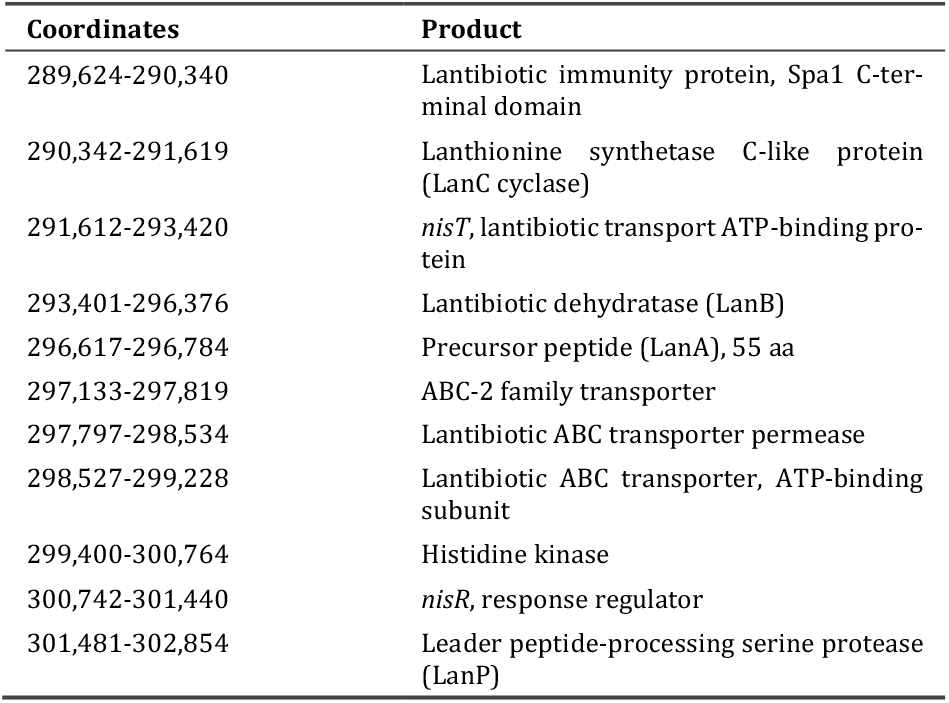

| Coordinates | Product |
|---|---|
| 289,624-290,340 | Lantibiotic immunity protein, Spa1 C-ter-minal domain |
| 290,342-291,619 | Lanthionine synthetase C-like protein (LanC cyclase) |
| 291,612-293,420 | *nisT*, lantibiotic transport ATP-binding pro-tein |
| 293,401-296,376 | Lantibiotic dehydratase (LanB) |
| 296,617-296,784 | Precursor peptide (LanA), 55 aa |
| 297,133-297,819 | ABC-2 family transporter |
| 297,797-298,534 | Lantibiotic ABC transporter permease |
| 298,527-299,228 | Lantibiotic ABC transporter, ATP-binding subunit |
| 299,400-300,764 | Histidine kinase |
| 300,742-301,440 | *nisR*, response regulator |
| 301,481-302,854 | Leader peptide-processing serine protease (LanP) |

The cluster is complete in the biosynthetic sense: a dehydratase and a cyclase to install the thioether rings, a dedicated transporter, a leader-cleaving protease, two-component regulation, and an immunity protein. Upstream of it, the flanking genes are a sigma-70 family factor at 288,952 and methionyl-tRNA synthetase at 286,564, both core chromosomal genes, which places the left boundary of the insertion.

The precursor encodes a 24-residue leader, MNNEDFNLDLVTISKENNSGASPR, and a 31-residue core, VTSKSLCTPGCKTGILMTCAIKTATCGCHFG. antiSMASH assigned it to class I with a RODEO score of 28 and predicted five thioether bridges, a monoisotopic mass of 2987.48 Da and a molecular weight of 2989.61 Da.

The core is nisin P. It matches the published mature nisin P peptide at all 31 residues, including the AIK hinge and the three-residue C-terminal truncation relative to nisin A that makes nisin P the shortest natural nisin variant. Nisin A is 34 residues with an NMK hinge, while nisin U and nisin P are both 31 residues with PLK and AIK hinges respectively; the alignment below sets the three cores side by side, with the nisin U sequence taken from BACTIBASE (29).

~~~
Nisin A   ITSISLCTPGCKTGALMGCNMKTATCHCSIHVSK
Nisin U   ITSKSLCTPGCKTGILMTCPLKTATCGCHFG
Nisin P   VTSKSLCTPGCKTGILMTCAIKTATCGCHFG
This work VTSKSLCTPGCKTGILMTCAIKTATCGCHFG
~~~

CompaRiPPson returned three antiSMASH-DB entries identical to the core across all 31 residues: NZ_FIKS01000005.1 in Streptococcus suis, and NZ_JADPLU010000018.1 and NZ_JAKVIE010000017.1 in Streptococcus pasteurianus. No S. agalactiae entry was returned. An identical 31-residue core in three species, with no synonymous divergence detectable at the peptide level, is more consistent with recent horizontal transfer than with vertical descent from a common ancestor. The chromosome carries both the peptide and a determinant of resistance to it: the nisin resistance protein annotated at 2,035,261 lies 1.7 Mb from this cluster, so production and resistance coexist in the isolate at separate loci.

### Predictions excluded, and annotation disagreements

antiSMASH predicted three regions; one is a genuine specialised-metabolite cluster. Region 1 (249,571-306,376) merged two unrelated loci roughly 30 kb apart: the lanthipeptide cluster above, and the *cyl* operon, classified as an arylpolyene. The *cyl* genes recovered were *cylX, cylD, cylG, cylZ, cylI, cylJ, cylK, cylF, cylA, cylB, cylE* and *acpC*. This is the granadaene biosynthetic operon, the beta-haemolysin/cytolysin and pigment locus present in essentially all GBS (30) and the basis of the chromogenic media used in maternal screening. It is reported here as an intact known virulence operon, not as a novel cluster, and serves as a positive control on the prediction.

Region 2 (418,300-439,173), called T3PKS, contains FemA, RecN, the arginine repressor, FolD, phosphoglucomutase and exodeoxyribonuclease VII. Region 3 (1,722,003-1,763,266), called terpene-precursor, contains mevalonate kinase, diphosphomevalonate decarboxylase, phosphomevalonate kinase, isopentenyl-diphosphate delta-isomerase and HMG-CoA reductase, that is, the mevalonate pathway. Both regions are primary metabolism, essential in Firmicutes, and neither is reported as a biosynthetic gene cluster. No lasso peptide, thiopeptide or sactipeptide region was predicted.

| **ST** | **n** | **Serotype(s)** |
|---|---|---|
| 1010 | 15 | IV |
| 17 | 12 | III |
| 1 | 11 | II (6), Ib (4), V (1) |
| 23 | 7 | Ia |
| 19 | 7 | V (5), III (2) |
| 498 | 7 | V (4), Ib (3) |
| 28 | 5 | II |
| 347 | 3 | II |
| 24 | 3 | V (2), II (1) |
| untypeable | 3 | Ia |
| 10 singletons | 10 | various |

Serotype frequencies are II 17 (20.5%), III 16 (19.3%), IV 15 (18.1%), V 14 (16.9%), Ia 12 (14.5%) and Ib 9 (10.8%). No genome was serotype VI to IX and none was non-typeable. GBS-SBG returned two serotype III subtype calls for 16 genomes, all within serotype III; these were collapsed to III, the unit used in the literature.

Two points follow from the distribution. The worldwide meta-analysis of maternal colonisation reports no serotype data at all for the Caribbean, while estimating the region’s colonisation prevalence at 34.7%, the highest of any region, from three countries and 1,137 women (31). These figures are among the first serotype data for the Caribbean, and they place serotype IV at 18.1%, concentrated entirely in ST1010, well above what is usual for a serotype that was uncommon until its recent emergence in adult invasive disease in North America (32). Separately, all six serotypes found here fall within the hexavalent GBS6 conjugate vaccine, which targets Ia, Ib, II, III, IV and V and is estimated to cover about 98% of isolates causing neonatal invasive disease (33). On serotype alone, this population is fully within that coverage.

#### A note on annotation disagreement

geNomad’s gene table and the Bakta annotation disagreed at four positions in these regions. Most consequentially, geNomad assigned the protein at 286,564-288,447 to a site-specific recombinase of the DNA invertase Pin family, whereas Bakta calls it methionyl-tRNA synthetase, an essential housekeeping gene; the length and context support Bakta. geNomad also assigned CONJScan conjugation profiles to proteins that Bakta annotates as a peptidylprolyl isomerase (313,160-313,843) and as an HTH Cro/C1 regulator and a replication initiation factor (1,112,360-1,113,592 and 1,134,832-1,135,860). Because geNomad’s marker set is built for viral and plasmid classification, its assignments outside that domain were not used; all element boundaries reported here derive from the Bakta annotation.

### The comparison collection: nineteen sequence types and six capsular serotypes

Eighty-three of the 93 assemblies from the Dominican maternal colonisation survey (19) were available for download, 82 at contig level and one complete. They resolve into 19 sequence types plus three untypeable, and six capsular serotypes. The frequencies below agree closely with those the source study reports for its full set of 93 isolates, which supports the reanalysis; what follows are the analyses that study did not perform.

Four sequence types carry more than one capsular type: ST1 appears as II, Ib and V across its eleven genomes, ST19 as V and III, ST498 as V and Ib, and ST24 as V and II. Capsular switching within a conserved genomic background is known in GBS, though reported as rare (34), and genomewide recombination across the species is well documented (35). Four apparent instances in one collection is relevant to vaccine coverage estimates, which are calculated per serotype (33).

### Prevalence of the nisin P operon

Twelve of the 83 genomes carry all seven cluster-specific genes. No genome carried a partial set: across 83 fragmented assemblies, a 13 kb locus is either wholly present or wholly absent, so the absences are not artefacts of assembly fragmentation. All twelve encode a precursor identical to the sequenced isolate’s at all 55 residues. Including the isolate, 13 of 84 genomes (15.5%) carry the operon. The previously published figure for GBS was one of thirteen faecal isolates.

| ST / serotype | With operon | Total |
|---|---|---|
| 19 / V | 5 | 5 |
| 347 / II | 3 | 3 |
| 28 / II | 2 | 5 |
| 498 / Ib | 1 | 3 |
| 2116 / II | 1 | 1 |
| 1010 / IV | 0 | 15 |
| 17 / III | 0 | 12 |
| 1 / II, Ib, V | 0 | 11 |

The operon occurs in five sequence types and is absent from the three largest lineages, which together account for 38 of the 83 genomes (Figure 2). Within ST28/II, two of five genomes carry it; within ST498/Ib, one of three; and within ST19, all five serotype V genomes carry it while neither serotype III genome does. Presence and absence within a single sequence type and capsular type excludes clonal inheritance as the explanation and identifies the locus as accessory.

**Figure 2.**
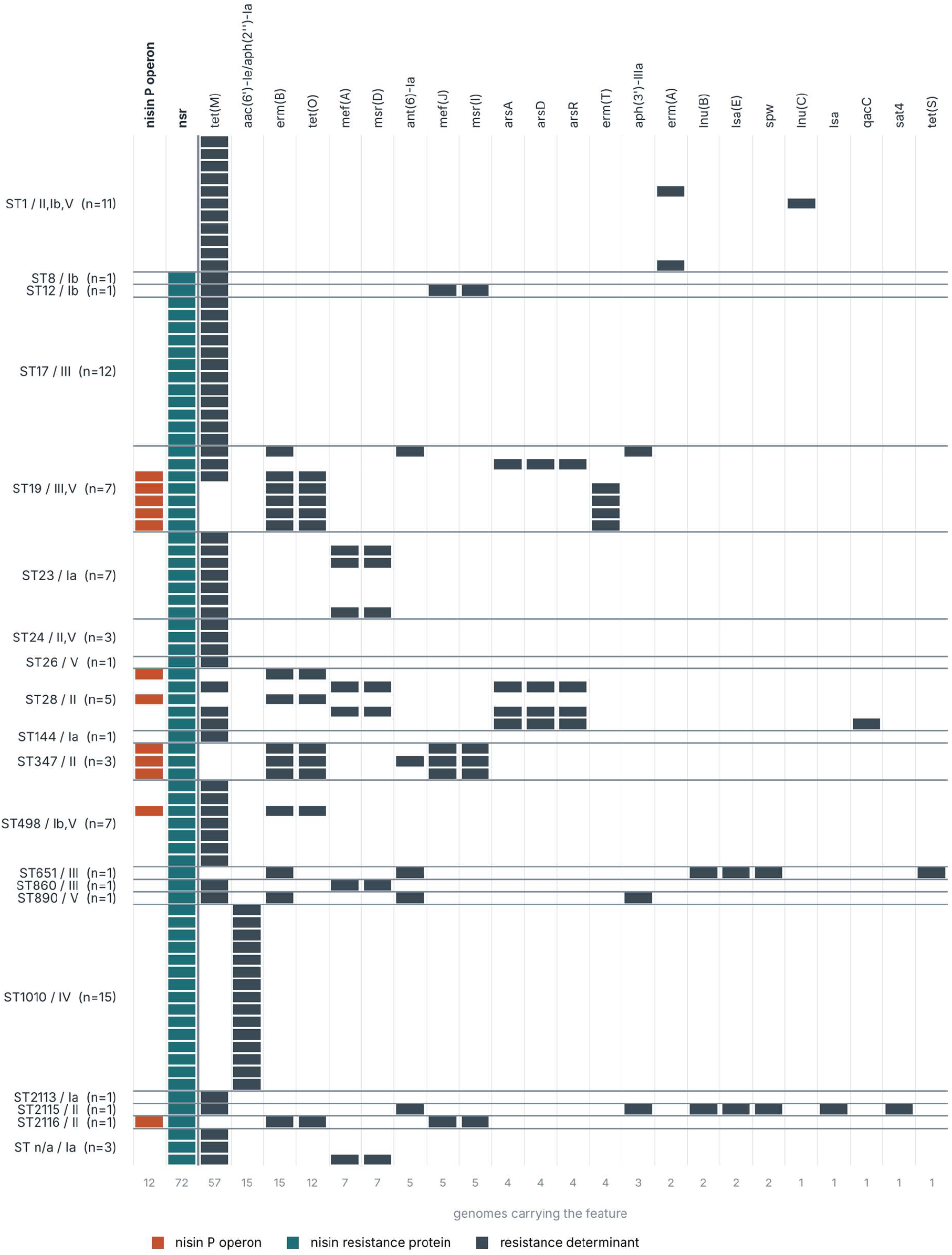
Lineage structure of accessory content across 83 Dominican group B Streptococcus genomes. Each row is a genome, grouped by sequence type; each column is a feature, ordered by frequency. The nisin P operon occurs in five sequence types and in none of the three largest lineages. The nisin resistance protein is present in every genome except the eleven of ST1. ST1010/IV carries a single determinant, aac(6′)-Ie/aph(2″)-Ia, in all fifteen genomes, and is the only lineage without tet(M). Every carrier of the operon also carries *erm(B)* and tet(O).

### *tet(O)* and *erm(B)* co-occur with the operon

In the closed genome the lanthipeptide cluster and the resistance element are 96 bp apart. Across the 83 draft genomes, the same linkage appears as co-occurrence.

|  | ***tet*(O) present** | ***tet*(O) absent** | ***erm*(B) present** | ***erm*(B) absent** |
|---|---|---|---|---|
| Operon present (12) | 12 | 0 | 12 | 0 |
| Operon absent (71) | 0 | 71 | 3 | 68 |
| Fisher, two-tailed | p = 1.03 x 10^-14 |  | p = 4.70 x 10^-12 |  |

Odds ratios after Haldane-Anscombe correction are 489 (95% CI 23.8 to 10,066) for *erm(B)* and 3,575 (95% CI 67.8 to 188,623) for *tet(O)*; both intervals are wide because one cell of each table is empty. For *tet(O)* the association is complete: every carrier of the operon carries *tet(O)* and no noncarrier does, giving *tet(O)* 100% sensitivity (95% CI 73.5and 100% specificity (95% CI 94.9-100) as a marker of the element in this population; the sensitivity interval is wide because it rests on twelve carriers. In the Dominican collection, *tet(O)* does not occur outside this element.

Twelve of the fifteen *erm*(B)-positive genomes (80%) carry the operon. Ten of the twelve carriers have *tet*(O) in place of *tet*(M) rather than in addition to it; only two carry both.

The two observations are independent of each other and of clonal structure. The twelve carriers span five sequence types, and within ST28/II and ST498/Ib the element is present in some genomes and absent from others, so the co-occurrence is not a single clone carrying two genes by descent. The physical linkage is established separately, in the closed assembly, where the two loci are contiguous.

| Genome | ST / serotype | Resistance genes |
|---|---|---|
| GCA_034335685.1 | 19 / III | *erm*(B), *tet*(M), *ant*(6)-Ia, *aph*(3')-IIIa |
| GCA_034335905.1 | 890 / V | *erm*(B), *tet*(M), *ant*(6)-Ia, *aph*(3')-IIIa |
| GCA_034336325.1 | 651 / III | *erm*(B), *tet*(S), *lsa*(E), *lnu*(B), *spw*, *ant*(6)-Ia |

The three erm(B)-positive genomes that lack the operon carry different elements. The first two carry erm(B) alongside tet(M) and two aminoglycoside determinants, the gene combination characteristic of the Tn916 family. The third carries a distinct multiresistance array built around lsa(E), lnu(B) and spw. At least three independent genetic routes to macrolide and lincosamide resistance therefore circulate in Dominican GBS, of which only one also carries a lantibiotic.

The second element moves independently of the first. mef(J) and msr(I) were found in five genomes: the three ST347/II, one ST2116/II, and GCA_034336725.1 (ST12/Ib), which carries mef(J), msr(I) and tet(M) but not the nisin P operon or tet(O). The two elements delimited in the closed genome are therefore separable in the population as well, and the sequenced isolate carries both.

### Nisin resistance and a susceptible lineage

The nisin resistance protein was found in 72 of 83 genomes, including all twelve carriers of the operon; no carrier lacked it, so production is invariably accompanied by a resistance determinant. Eleven genomes carry neither the operon nor the resistance protein, and all eleven are ST1: the entire ST1 lineage of the collection, across all three of its capsular types. The division is exactly by lineage and admits no exception; every genome outside ST1 carries the resistance protein, and no ST1 genome does. Absence of the resistance protein does not by itself establish phenotypic susceptibility, since other mechanisms exist, but it makes a lineagewide prediction that can be tested directly.

One genome, GCA_034336325.1 (ST651, serotype III), returned a hit to the immunity protein without any other cluster gene. A single tblastn hit in a draft assembly is not sufficient evidence of an acquired immunity determinant, and this observation requires confirmation of identity and genomic context before it is interpreted.

### The population resistome

Every one of the 83 genomes carried at least one acquired resistance determinant. By class, 68 of 83 (81.9%) carry a tetracycline determinant, 27 of 83 (32.5%) a macrolide or lincosamide determinant, and 20 of 83 (24.1%) an aminoglycoside determinant. Fifty-four genomes carry determinants of one class, 13 of two, 11 of three, three of four and two of five.

The macrolide and lincosamide burden is carried by five distinct determinants rather than one dominant element: *erm(B)* in 15 genomes, *mef(A)* with *msr(D)* in 7, *mef(J)* with *msr(I)* in 5, *erm(T)* in 4 and *erm(A)* in 2, with *lsa(E)* and *lnu(B)* accounting for a further small number. One genome, GCA_034336745.1 (ST2115/II), additionally carries two full-length hits that AMRFinderPlus assigns to the Lsa family rather than to a known allele: both cover 100% of the 492-residue reference at 90.0% identity to Lsa(C), on two separate contigs. These are divergent Lsa-family ribosomal protection proteins, not truncated or partial genes, and they are the only resistance determinants in this collection that could not be resolved to an allele. *mef(A)* and *msr(D)* form the efflux pair carried by the MEGA element (36), while *tet(M)* and *tet(O)* both encode ribosomal protection proteins (37). For a single-country collection this diversity is itself the finding: no one element dominates, and interventions aimed at a single mechanism would not reach most carriers. *erm(T)* is confined to four of the five ST19/V carriers of the nisin P element, and whether it lies inside that element or adjacent to it is unresolved in draft assemblies. Accessory content can diverge sharply within a single sequence type. The five ST28/II genomes separate cleanly by accessory content: the three without the nisin P element carry tet(M) with arsRDA arsenic resistance and either mef(A)/msr(D) or qacC, while the two with the element carry erm(B) and tet(O) and nothing else. Same sequence type, same capsular type, mutually exclusive accessory repertoires. qacC encodes a quaternary ammonium compound efflux pump, which is relevant where chlorhexidine is used in obstetric antisepsis.

The two most resistant genomes in the collection are GCA_034336745.1 (ST2115/II), with eight determinants across five classes, and GCA_034336325.1 (ST651/III), with six.

### ST1010/IV carries high-level gentamicin resistance clonally

All fifteen ST1010/IV genomes carry *aac*(6’)-Ie/*aph*(2’’)-Ia, the bifunctional aminoglycoside-modifying enzyme, and no other acquired determinant. ST1010 is the largest lineage in the collection at 18.1%, and it is the only lineage in which *tet*(M) is absent; its resistome is a single gene, identical across all fifteen genomes.

This enzyme confers high-level gentamicin resistance, for which laboratory breakpoints are not yet harmonised (38), and abolishes the synergy between a beta-lactam and gentamicin, the combination used for GBS neonatal sepsis and meningitis (39) and for invasive disease in adults, although the role of aminoglycoside synergy has itself been questioned in endocarditis (40).

The association is not new. Creti and colleagues described high-level gentamicin resistance in serotype IV ST1010/CC452 in Italy, carried on a novel integrative conjugative element of about 45 kb, ICESag9931, derived from ICE_515_tRNA-Lys with a truncated Tn*3706* inserted into it (41). That study searched public databases for the lineage, listed Dominican ST1010 strains from 2021 among its matches, and did not determine their resistance status. The present reanalysis answers that question: the determinant is present in 15 of 15 Dominican ST1010/IV genomes.

Two things follow. Whether the gene sits on ICESag9931 in these genomes is unresolved and testable against the published element. And the phenotype should be confirmed by high-level gentamicin screening on available isolates, because a determinant that is uniform across the country’s most common colonising lineage bears on local empirical regimens for invasive disease.

### CC17 carries no macrolide determinant

All twelve ST17/III genomes carry *tet*(M) and nothing else. ST17 is the hypervirulent clonal complex associated with late-onset neonatal disease (42), and in this collection it has no macrolide or lincosamide determinant at all. Clindamycin prophylaxis would therefore be expected to remain effective against the lineage of greatest neonatal concern, which is worth stating alongside the resistance findings.

## Discussion

The central observation is that in this population clindamycin resistance is genetically coupled to a trait that is plausibly advantageous during colonisation. An element that carries both can be maintained by either selective pressure. Macrolide use in the community would select for carriers and thereby spread a lantibiotic producer; competition during vaginal or intestinal colonisation would select for carriers and thereby spread *erm*(B). Bacteriocin production is known to augment niche competition in vivo (43, 44), and in enterococci it has been shown to facilitate the nosocomial emergence of a vancomycin-resistant lineage (45), which is the same coupling proposed here in a different organism. Co-selection of this kind is well described for metal and biocide resistance linked to antibiotic resistance, but the linked trait here is not a second resistance; it is an antibacterial weapon. If the coupling holds, reducing macrolide pressure would not by itself be expected to reduce the frequency of *erm*(B) in Dominican GBS, because the element has a second reason to persist.

The argument rests on two independent lines of evidence that are not substitutes for each other. The closed assembly shows the two loci 96 bp apart, which drafts cannot show; the population screen shows them co-occurring at p = 10^-14 across five sequence types with within-type variation, which a single genome cannot show. Neither alone would be persuasive. This is the specific contribution of closing one genome from a collection that otherwise consists of drafts: it converts a statistical association into a described structure.

Nisin P had been reported from one *S. agalactiae* isolate, a human faecal strain in which the operon was present in one of thirteen sequenced stool isolates from the same study (17). At 15.5% across 84 Dominican genomes, the operon is substantially more frequent here, though the comparison is loose: both sets are colonising isolates, but from different body sites, countries and sampling frames. The identity of the 31-residue core across *S. agalactiae, S. pasteurianus* and *S. suis*, with the cluster first described in the latter two, points to an interspecies reservoir, most plausibly in the gut, and to acquisition by GBS rather than origin within it. Genomic surveys of GBS elsewhere in Latin America, such as the recent Argentine collection (46), are the obvious comparison set for establishing whether this element is regional or more widely distributed.

The absence of the operon from the three largest lineages, and its complete absence from ST1 together with the absence of the nisin resistance protein from that same lineage, describes a population with a structured competitive landscape rather than a uniform one. Whether that structure is a cause or a consequence of the element’s distribution is not answerable from genomes. The deferred-antagonism assay proposed below is the shortest route to an answer: if ST1 isolates are inhibited by the producer and the 60 resistance-protein-positive non-producers are not, the genotype predicts the interaction, and the element’s persistence has a measurable fitness basis.

Two further findings bear on clinical practice without belonging to that argument. Surveillance that reports only the resistance gene would record the three erm(B) routes described here as one phenomenon, when they have different mobility, different cargo and probably different drivers, and only one is linked to a lantibiotic. And the clonal fixation of aac(6’)-Ie/aph(2’’)-Ia in all fifteen ST1010/IV genomes carries more immediate weight than the composite element: beta-lactam and gentamicin synergy underpins the treatment of invasive GBS disease, high-level gentamicin resistance abolishes it, and the determinant is uniform across the most common colonising lineage in the collection. That extends a published observation (41) rather than making a new one, and it warrants phenotypic confirmation before it bears on any regimen. The converse finding is reassuring and equally worth reporting: CC17, the hypervirulent lineage, carries no macrolide or lincosamide determinant here. Finally, the predicted inducible MLSb phenotype of the sequenced isolate is the point where this work becomes actionable for a single patient rather than a population. An intact attenuator upstream of *erm*(B) predicts that clindamycin may test susceptible and fail in vivo. A D-test costs a plate.

### Limitations

#### The comparison set is one collection of colonising isolates

All 83 genomes come from a single study: rectovaginal swabs from 350 near-term pregnant women at one tertiary hospital in Santo Domingo between September 2021 and March 2022 (19). They are colonising isolates from one city over seven months, not a national sample and not invasive disease isolates. Prevalence figures describe that collection. Ten of the study’s 93 sequenced isolates had no assembly available for download and are missing from this reanalysis. No clinical, demographic or temporal metadata beyond what the source study reports were available for individual genomes.

#### Co-occurrence is not linkage

Physical contiguity of the lanthipeptide cluster and the resistance element is demonstrated only in the closed genome. In the 83 drafts the relationship is statistical. A genome in which the two loci cooccur but lie apart would be indistinguishable in a draft assembly, and we cannot exclude that the association in some carriers reflects two elements acquired together rather than one.

#### No phenotype was measured

The inducible MLSb prediction, the activity of the peptide, the susceptibility of ST1, the high-level gentamicin resistance of ST1010 and the transmissibility of either element are all inferences from sequence. None has been tested. No antibiogram for the sequenced isolate was available at the time of writing.

#### The peptide was not isolated

Identification as nisin P rests on identity of the predicted core sequence to the published mature peptide, and on CompaRiPPson matches. No mass spectrometry was performed, so neither production nor the predicted 2987.48 Da species has been confirmed, and post-translational modification is predicted rather than observed.

#### Element boundaries are inferred

Both elements are delimited from the outermost annotated mobility and conjugation genes flanking their cargo. Attachment sites, direct repeats and excision were not determined experimentally, and the figures of 34.4 kb and 35 kb should be read as approximate. The tRNA-Lys adjacency fixes one boundary of the second element more confidently than annotation alone.

#### Presence calls depend on thresholds

The operon criterion excludes three genes that cross-hit native streptococcal transporters and regulators; a different threshold or a different gene set would give a different count. The criterion is stated in the methods so that it can be reproduced or disputed.

#### One observation rests on a single hit

The immunity protein found in GCA_034336325.1 without the rest of the cluster is a single tblastn hit in a draft assembly and is reported as unverified.

#### No viable stock, and no methylome

The isolates were discarded after DNA extraction, so no living culture of this organism exists and every phenotypic test proposed below would require re-isolation of the lineage from a new specimen. Raw signal data were not retained either, so base modification analysis is not possible. Any future methylome work on this lineage requires resequencing.

#### The two libraries are technical replicates

They quantify the reproducibility of library preparation, sequencing and basecalling. Both derive from colony sweeps of the same specimen, so they exclude a fixed-ratio two-clone mixture at the five discordant positions but say nothing about within-host diversity elsewhere in the genome, and the isolate represents one patient.

### Experiments this work sets up

Each of these tests a specific prediction made above, and each is feasible with standard clinical microbiology equipment.

1. **Double-disc diffusion (D) test on a re-isolated ST347/II strain carrying *erm*(B)**. Tests whether the intact attenuator upstream of *erm*(B) produces an inducible rather than constitutive MLSb phenotype. A positive D-test means clindamycin may report susceptible and fail in vivo, which bears directly on intrapartum prophylaxis in women with reported penicillin allergy.
2. **Deferred-antagonism assay of an operon carrier against ST1 isolates**. Tests the lineage-wide prediction that ST1, which lacks both the operon and the nisin resistance protein, is inhibited by a producer, while non-producers carrying the resistance protein are not. This is the single experiment that would give the element’s persistence a measured fitness basis.
3. **Mass spectrometry of culture supernatant from a carrier**. Looks for the predicted 2987.48 Da monoisotopic species, confirming that the operon is expressed and the peptide modified as predicted. Without it, production remains an inference from gene content.
4. **Filter-mating conjugation assay**. Tests whether element 1 transfers as a unit, carrying the nisin P operon together with *erm*(B) and *tet*(O), using *tet*(O) as the selectable marker and a recipient lacking it. A positive result would convert the composite element from a described structure into a demonstrated vehicle.
5. **High-level gentamicin screening on ST1010/IV isolates**. Confirms phenotypically the determinant found clonally in all fifteen genomes of the largest lineage, and establishes whether betalactam and gentamicin synergy is compromised in it.
6. **Single-colony re-sequencing of a carrier**. Both libraries here derive from colony sweeps, and the allele-fraction test excludes a fixed-ratio two-clone mixture at the discordant positions. Sequencing single-colony picks from a fresh plate would extend that test to the whole genome and establish whether carriers of this lineage harbour more than one clone at all.

## Acknowledgements

The authors thank the originating diagnostic laboratory, which asked not to be identified for providing the bacterial isolate, and the Instituto Especializado de Estudios Superiores Loyola and its Department of Research for access to the bioscience laboratories.

## Author contributions

L.G.E. took part in every stage of the work, from isolate handling through to bioinformatic analysis, manuscript writing, and data analysis. D.S.D.L.C.-C. performed the DNA extraction. H.B.L.C. conceived and supervised the study, commissioned the sequencing, performed the assembly, annotation, comparative genomic analysis and statistics, and took part in manuscript writing. All authors read and approved the final version.

## Materials and Methods

### Isolate, sequencing and ethics

The isolates were recovered in the course of routine diagnosis at a private clinical diagnostic laboratory in the Dominican Republic, which asked not to be identified, from a vaginal swab submitted for analysis. Once the diagnostic result had been issued, they formed part of residual material scheduled for disposal, from which they were selected and transferred to the authors. No clinical or demographic information accompanied them, and the diagnostic care of the source patient was complete and unaffected.

Two blood agar plates from this specimen were processed. All colonies on each plate were harvested as a sweep, and DNA was extracted separately from each, giving two independent cultures, extractions and libraries, designated S1 and S2. The originating laboratory confirmed that both plates derive from the same specimen. No viable stock was retained; the remaining material was discarded after extraction.

The sequenced isolate and the comparison genomes therefore come from different sampling frames. The comparison genomes were collected prospectively by screening asymptomatic near-term pregnant women; the sequenced isolate comes from a specimen submitted to a diagnostic laboratory. Whether the source patient was symptomatic is not documented, and no inference about that is drawn here.

Phenotypic susceptibility testing was not available from the originating laboratory, and no viable stock of the isolate was retained, so no phenotype can be obtained from it. Every phenotypic test proposed in the discussion requires re-isolation of the lineage from a new specimen.

Read statistics were computed with nanoq (47). The two libraries are not equivalent in yield: S1 gave 915,370 reads totalling 5.95 Gb with read N50 7,673 bp and mean quality Q23.71, while S2 gave 1,248,652 reads totalling 7.45 Gb with read N50 7,281 bp, mean read length 5,965 bp and mean quality Q23.44. S2 therefore carries 36% more reads and 25% more bases at slightly shorter read length and slightly lower quality.

Library preparation and sequencing were performed under contract by Psomagen (Rockville, MD, USA). The provider supplied basecalled reads only: the library preparation kit, the flow cell product code and the barcode assignments were not available to the authors, so the account of the two libraries given above rests entirely on the metadata carried in the read headers. The 2,826 distinct channels exclude a MinION or Flongle flow cell, which carry 512 and 126 channels, and indicate a PromethION-class flow cell, but the specific product was not confirmed. Basecalling used the Oxford Nanopore model <u>dna_r10.4.1_e8.2_400bps_sup@v5.2.0,</u> that is, R10.4.1 chemistry with the E8.2 pore at 400 bases per second and the super-accuracy model, run on an NVIDIA RTX 6000 Ada GPU; the sequencing run is dated 17 June 2026.

Reads were filtered with Filtlong (48) using `--min_length 1000 --keep_percent 90 --target_bases 420000000`, a target of approximately 203-fold coverage of the 2.07 Mb chromosome. Filtering retained 26,710 reads totalling 420,010,410 bases for S1 (read N50 15,160 bp, mean quality Q25.1) and 26,266 reads totalling 420,011,876 bases for S2 (read N50 15,452 bp, mean quality Q25.2); read N50 roughly doubled and mean quality rose by about 1.5 Phred units in both. Assembly used Flye (49) with `--nano-hq`, the mode intended for R10.4.1 chemistry with super-accuracy basecalling. Long-read-only reconstruction of bacterial genomes with R10.4.1 has been evaluated and found accurate enough to forgo short-read polishing (50). Consensus assembly across several assemblers (51) was not used: the two libraries were assembled independently and the two assemblies compared, which tests reproducibility rather than smoothing it away.

### Two libraries from one specimen, and the replicate comparison

The two read sets were sequenced simultaneously and basecalled together. Their records share one read group, one device identifier and one run start time; they draw on the same 2,826 channels, of which 2,825 are common to both; and they span the same acquisition window, beginning 438 ms apart and ending 233 ms apart across a run of 4 h 44 min. They share no read identifiers. Neither a positional nor a temporal partition of a single read set is compatible with this.

Despite that complete interleaving, the two sets differ measurably in composition: read N50 7,673 bp against 7,281 bp, a difference of 392 bp or 5.1%, and mean quality Q23.7 against Q23.4.

To establish whether a difference of that size can arise from partitioning a single pool, the two sets were combined and three independent random partitions of the same proportion were drawn from the combined reads. These gave read N50 of 7,463, 7,461 and 7,457 bp, a total spread of 6 bp, with mean quality identical to one decimal place in all three. The observed difference between S1 and S2 is approximately 65 times that spread (Figure 3). Partition of a single read set is therefore excluded empirically, not only by the positional and temporal evidence above.

**Figure 3.**
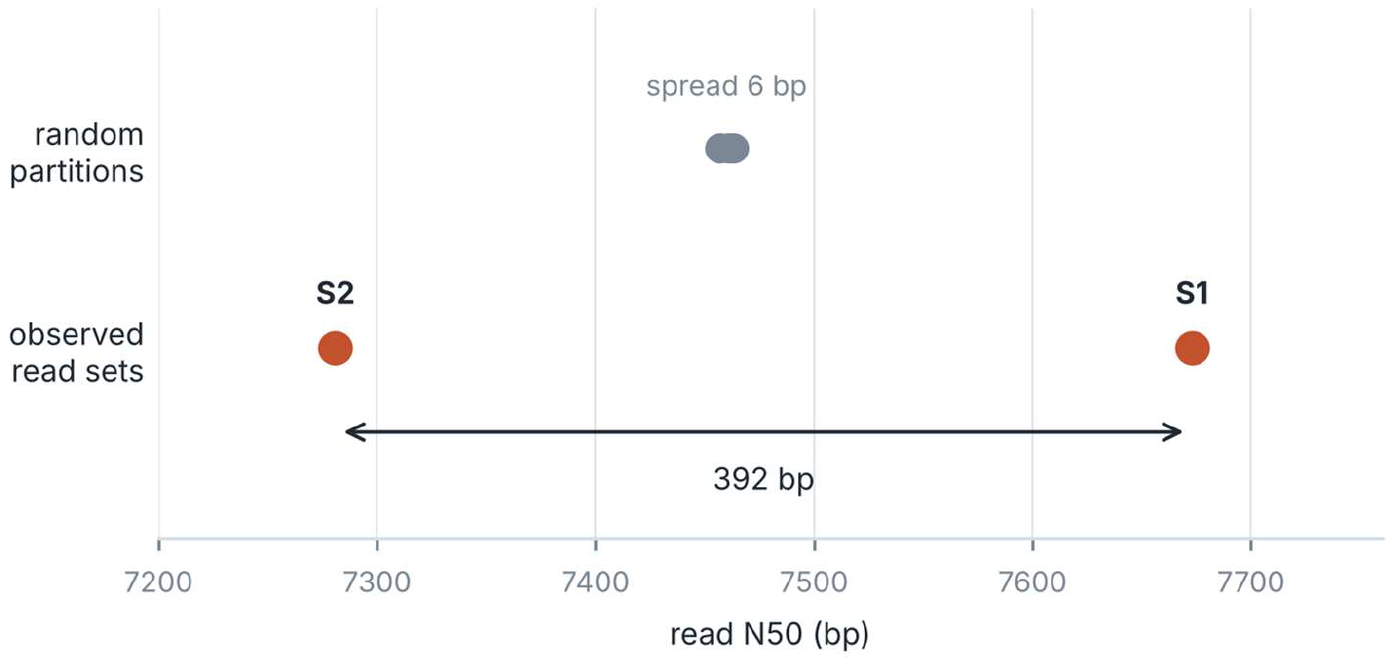
The two read sets are not a partition of one. Three random partitions of the pooled reads, drawn at the same proportion with independent seeds, give read N50 values spanning 6 bp. The observed sets S1 and S2 differ by 392 bp, approximately 65 times that spread, which excludes partition of a single read set.

S1 and S2 are distinct populations of molecules sequenced together on one flow cell, which is what two separately prepared libraries multiplexed in one run would produce. No barcode tag is retained in the delivered FASTQ files, so the demultiplexing step itself could not be recovered from the data, and the inference rests on the evidence set out here.

The comparison between them therefore measures the combined reproducibility of library preparation, sequencing and basecalling, not basecalling alone. Reads from each library were mapped to the S1 assembly with minimap2 and allele counts at the five discordant positions were taken from `samtools mpileup` without mapping-quality or base-quality filtering; the script that reproduces this is included in the code repository.

The two assemblies were aligned with `dnadiff` from the MUMmer package (52). Each discordant position was then examined at the read level: both read sets were mapped to the S1 assembly with minimap2 (53) and base composition at the five positions was tabulated from `samtools mpileup` (54) output, so that a genuine difference between replicates could be distinguished from a position where both read sets are internally ambiguous. Minor-allele fractions were then compared across the three substituted positions to test whether a single mixing ratio, as a mixture of two clones would produce, could account for them.

At the three substituted positions both read sets carried both bases at depths of 172 to 228 reads against a genome-wide mean near 200, so the ambiguity is present in each replicate independently and at normal depth rather than reflecting a collapsed repeat that one assembly resolved differently. Because both libraries derive from sweeps of all colonies on a plate rather than from single-colony picks, within-host clonal diversity was a competing explanation, and read-level phasing cannot adjudicate it: the three substitutions lie 260 kb, 596 kb and 856 kb apart and no read spans any pair. The mixing-ratio test is therefore the decisive one. A mixture of two clones has a single ratio, so every site that distinguishes them must show the same minor-allele fraction. Pooling both libraries, the alternative base accounted for 29.4% of reads (95% CI 25.2-34.0) at position 1,198,656, 46.6% (41.1-52.2) at 1,794,246 and 55.8% (50.6-61.0) at 2,054,407, with the same rank order in each library separately (24.7%, 44.0% and 55.5% in S1; 34.3%, 49.4% and 56.1% in S2). Two independent colony sweeps would not be expected to share a mixing ratio, still less to reproduce the same ordering of fractions across sites, whereas context-dependent basecalling error is deterministic given the local sequence and recurs in both runs. Position 1,198,656 further yielded five distinct observations (G, C, T, A and a deletion), which a biallelic difference between two clones cannot produce. The mechanism by which the two assemblies come to differ is directly visible at position 2,054,407, where the S1 assembly called A although 55.5% of its own reads carried G, and the S2 assembly called G: the difference between the genomes at that site is the consensus step resolving a near-even position to opposite bases in the two runs.

To test whether the two read sets could instead be a partition of one, they were concatenated and three random partitions of the same proportion (42.3%) were drawn with independent seeds, each read retained with that probability. Read N50 and mean quality were computed for each partition with nanoq and compared with the observed values for S1 and S2.

### Annotation and typing

The closed assembly was annotated with Bakta 1.12.2 (55) against database v6.0 (light, 2025-02-24), with `--genus Streptococcus --species agalactiae`. Multilocus sequence typing used the `sagalactiae` scheme (*adhP, pheS, atr, glnA, sdhA, glcK, tkt*) (56), and the clonal complex was assigned from the PubMLST profile (57). Capsular serotype was called with GBS-SBG (58) from the assembly.

### Resistance determinants

AMRFinderPlus (59) was run on nucleotide input with `--organism Streptococcus_agalactiae –plus`, twice on the closed genome: version 3.12.8 with the 2024 database and version 4.2.7 with database 2026-08-07.1. Calls were compared to those produced independently by Bakta’s own AMRFinder expert system. AMRFinderPlus 4.2.7 with database 2026-08-07.1 was then run on all 83 comparison genomes; identical duplicate rows in its output were collapsed on the key accession, element symbol, contig and start coordinate.

### Mobile elements and biosynthetic gene clusters

Plasmids and proviruses were predicted with geNomad 1.13.0 (60) `end-to-end` against database v1.9. Mobility and conjugation genes were taken from the Bakta annotation and cross-checked against geNomad’s gene table; where the two disagreed, the Bakta assignment was used, and the disagreements are reported. Element boundaries are inferred from the outermost mobility or conjugation gene flanking each cargo region and are therefore approximate.

Biosynthetic gene clusters were predicted with antiSMASH 8.0.4 (61) on the Bakta GenBank output using `--taxon bacteria --genefinding-tool none --cb-general --cb-knownclusters --cb-subclusters --asf --pfam2go --smcog-trees`. Predicted regions were curated manually against their gene content; regions whose genes were entirely primary metabolism or a previously characterised virulence operon were excluded, and the exclusions are reported. The lanthipeptide core was compared to published RiPP cores with CompaR-iPPson against antiSMASH-DB 4.0 and MIBiG 4.0.

| Tool | Version | Database |
|---|---|---|
| nanoq | 0.10.0 | - |
| Filtlong | 0.3.1 | - |
| Flye | 2.9.6 | - |
| minimap2 | 2.31 | - |
| SAMtools | 1.24 | - |
| MUMmer / dnadiff | 4.0.1 (dnadiff 1.3) | - |
| Bakta | 1.12.2 | v6.0 light, 2025-02-24 |
| mlst | 2.35.0 (bioconda build hdfd78af_1) | PubMLST *sagalactiae* |
| GBS-SBG | commit 9e53847 | bundled *cps* reference set |
| AMRFinderPlus | 3.12.8 and 4.2.7 | 2024-07-22.1 and 2026-08-07.1 |
| geNomad | 1.13.0 | v1.9 |
| antiSMASH | 8.0.4 | ASDB 4.0, MIBiG 4.0, Pfam-A 35.0 |
| BLAST+ | 2.16.0 | - |
| SciPy | 1.17.1 | - |

### Software versions

Analyses were run in separate conda environments; BLAST+ 2.16.0 is the version in the environment from which the comparative tblastn screens were executed, and other environments carry 2.17.0. All versions are as recorded in the project conda environments at the time of analysis.

| **Record** | **Count** | **Identifier** |
|---|---|---|
| BioProject | 1 | PRJNA1552734 |

The five positions at which the two replicates differ are reported in a supplementary table with their read-level base composition, flagged as base-caller-ambiguous rather than resolved to one state.

### Comparison genomes

The 83 Dominican *S. agalactiae* assemblies used for comparison are public; their accessions, sequence types, serotypes, element status and resistance determinants are listed in Supplementary Table S1, generated as `tabla_final_dr.tsv`

### Comparative screen

Assemblies deposited from the Dominican maternal colonisation survey of Laycock et al. (19), BioProject PRJNA945321, were retrieved with the NCBI `atasets` command-line tool, giving 83 genomes (82 contig-level, one complete). The source study reports 93 sequenced isolates; 83 assemblies were available for download at the time of analysis, and the ten absent isolates are a limitation of this reanalysis rather than of the original work. These are colonising isolates from near-term pregnant women, not invasive or clinical isolates. Query proteins were taken from the closed genome’s Bakta annotation and searched against a concatenated BLAST database of all 83 assemblies with tblastn (62).

Presence of the lanthipeptide operon was defined by the seven cluster-specific genes: the LanC cyclase, the LanB dehydratase, the LanA precursor, the two ABC transport components, the histidine kinase and the leader-processing protease. The *nisT* transporter, the *nisR* response regulator and the ABC ATP-binding subunit were excluded from the presence criterion because each returned hits in 81 to 83 genomes, reflecting homology with native streptococcal ABC transporters and two-component systems rather than the cluster. Thresholds were e-value 1 x 10^-10 and query coverage at least 70% for cluster genes, and e-value 1 x 10^-20 with at least 95% identity and 90% query coverage for resistance genes.

### Statistics

Two-by-two associations were tested with Fisher’s exact test, two-tailed, as implemented in SciPy (63). Association tests were computed from the presence and absence tables generated by the screen; the script that reproduces them from ‘operonP_dr.tsv’, ‘amr_dr.tsv’ and ‘st_serotipo_dr.tsv’ is included in the code repository. For the table containing a zero cell, the odds ratio and its 95% confidence interval were computed after a Haldane-Anscombe correction of 0.5 added to every cell. Minor-allele fractions at the positions discordant between the two libraries were compared across positions with a chi-square test of homogeneity on the two-by-three table of major and minor allele counts, and the 95% confidence interval of each fraction was obtained from an exact binomial test, both as implemented in SciPy.

## Data availability, deposition and ethics

### Deposition plan

The two libraries are one organism, so they are deposited as one BioSample with two sequencing runs. Depositing them as two isolates would place a technical replicate in the public record as two genomes.

### Code

All commands used for assembly, annotation, typing, the comparative screen and the statistics will be deposited in a public repository https://github.com/hanseln3s5/gbs-st347-nisinP.git

### Ethics

The isolate was recovered in routine diagnostic work at a private clinical diagnostic laboratory in the Dominican Republic, which asked not to be identified, and formed part of residual material scheduled for disposal once the diagnostic result had been issued. It was transferred to the authors under a release document issued by that laboratory, without any accompanying clinical or demographic information and with no means of linking it to a patient. All laboratory and computational work was carried out at the Instituto Especializado de Estudios Superiores Loyola, under its institutional responsibility. No identifiable patient information was transferred to the authors or is reported here, and no patient’s diagnostic care was affected.

