## Supplemental information 1 for "A closed genome of *Streptococcus agalactiae* ST347/II reveals a composite integrative element coupling nisin P biosynthesis to *erm(B)* and tet(O) in Dominican group B *Streptococcus*"

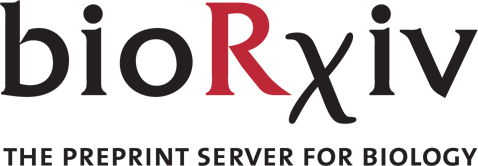

Supplementary Information for: A closed genome of Streptococcus agalactiae ST347/II reveals a composite integrative element coupling nisin P biosynthesis to erm(B) and tet(O) in Dominican group B Streptococcus

Leslys García-Encarnación, Dolinsky S. De La Cruz-Caminero, Hansel B. López-Chevalier*

### Contents

Supplementary Tables S1 to S11 are provided as a separate Excel workbook (Supplementary_Tables_S1-S11.xlsx). The README sheet of that workbook lists the sheets and the decisions that shaped them.

**Supplementary Table S1.** The 83 Dominican group B Streptococcus genomes analysed, with sequence type, capsular serotype, nisin P operon status, nisin resistance protein and acquired resistance determinants.

**Supplementary Table S2.** Nisin P operon by sequence type and serotype.

**Supplementary Table S3.** Frequency of acquired resistance and stress determinants across the comparison genomes.

**Supplementary Table S4.** tblastn hits per query protein, and the three proteins excluded from the presence criterion.

**Supplementary Table S5.** Association tests for tet(O) and erm(B) against operon presence, and the random-partition control for the two read sets.

**Supplementary Table S6.** Read and assembly statistics for both read sets, before and after filtering.

**Supplementary Table S7.** Gene inventory of element 1, which carries erm(B) and tet(O).

**Supplementary Table S8.** Gene inventory of element 2, which carries mef(J) and msr(I).

**Supplementary Table S9.** The eleven-gene nisin P biosynthetic cluster.

**Supplementary Table S10.** Software, databases and parameters.

Supplementary Table S11. Allele fractions at the three positions discordant between the two libraries, with confidence intervals and the test of homogeneity.

**Supplementary Note 1.** Analysis commands.

**Supplementary Note 2.** Decisions that shape the results, and what they exclude.

### Supplementary Note 1: Analysis commands

Commands are given in the order run. Paths are those of the analysis system. The complete scripts are deposited in the code repository cited in the data availability statement.

Read filtering and assembly

```
nanoq -i S1.fastq -s -vv
filtlong --min_length 1000 --keep_percent 90 --target_bases 420000000 \
  S1.fastq | gzip > filt/S1.filt.fastq.gz
flye --nano-hq filt/S1.filt.fastq.gz --out-dir flye/S1 --threads 16
```

Replicate comparison and the partition control

```
dnadiff flye/S1/assembly.fasta flye/S2/assembly.fasta -p s1s2
show-snps -ClrTH s1s2.delta
minimap2 -ax map-ont -t 16 flye/S1/assembly.fasta filt/S1.filt.fastq.gz \
  | samtools sort -@ 8 -o S1.bam - && samtools index S1.bam
minimap2 -ax map-ont -t 16 flye/S1/assembly.fasta filt/S2.filt.fastq.gz \
  | samtools sort -@ 8 -o S2_vs_S1.bam - && samtools index S2_vs_S1.bam
python3 fracciones.py flye/S1/assembly.fasta S1.bam S2_vs_S1.bam

for i in 1 2 3; do
  cat S1.fastq S2.fastq \
    | awk -v s=$i 'BEGIN{srand(s)} NR%4==1{k=(rand()<0.423)} k' \
    | nanoq -s
done
```

Annotation, typing and resistance

```
bakta --db /mnt/das2/db/bakta/db-light --output bakta_S1 --prefix S1 \
  --genus Streptococcus --species agalactiae --locus-tag S1 --threads 16 \
  flye/S1/assembly.fasta

mlst flye/S1/assembly.fasta
perl GBS-SBG/GBS-SBG.pl flye/S1/assembly.fasta

amrfinder -n flye/S1/assembly.fasta --organism Streptococcus_agalactiae --plus
# la base 2026-08-07.1 vive en el entorno amrfinder, no en gbs-annot:
export AMRFINDER_DB=$HOME/micromamba/envs/amrfinder/share/amrfinderplus/data/2026-08-07.1
```

Mobile elements and biosynthetic gene clusters

```
genomad end-to-end flye/S1/assembly.fasta genomad_S1 /mnt/das1/db/genomad_db

antismash --taxon bacteria --genefinding-tool none \
  --cb-general --cb-knownclusters --cb-subclusters \
  --asf --pfam2go --smcog-trees \
  --output-dir antismash_S1 --cpus 16 bakta_S1/S1.gbff
```

Comparative screen

```
datasets download genome accession --inputfile acc_dr.txt
for f in dr_genomes/ncbi_dataset/data/*/*.fna; do
  acc=$(basename "$(dirname "$f")")
  awk -v a="$acc" '/^>/{sub(/^>/,">"a"|")} {print}' "$f"
done > blastdb/dr_all.fna
makeblastdb -in blastdb/dr_all.fna -dbtype nucl -out blastdb/dr_all

tblastn -query clusterP_prot.faa -db blastdb/dr_all -evalue 1e-10 \
  -num_threads 16 -outfmt '6 qseqid sseqid pident qcovhsp' \
  | awk -F'\t' '$4>=70 {split($2,a,"|"); print $1"\t"a[1]}' | sort -u > operonP_dr.tsv

tblastn -query amr_query.faa -db blastdb/dr_all -evalue 1e-20 \
  -num_threads 16 -outfmt '6 qseqid sseqid pident qcovhsp' \
  | awk -F'\t' '$3>=95 && $4>=90 {split($2,a,"|"); print $1"\t"a[1]}' | sort -u > amr_dr.tsv
```

### Supplementary Note 2: Decisions that shape the results

Which genes define operon presence

Presence was called on seven cluster-specific genes: the LanC cyclase, the LanB dehydratase, the LanA precursor, two ABC transport components, the histidine kinase and the leader-processing protease. The nisT transporter, the nisR response regulator and the ABC ATP-binding subunit were excluded because each returns hits in 81 to 83 of the 83 genomes, reflecting homology with native streptococcal transporters and two-component systems rather than the cluster. Applying the criterion to all eleven genes would call every genome positive; applying it to the seven gives twelve. No genome carried a partial set, so the distinction does not rest on a threshold.

Serotype III subtypes

GBS-SBG returned two serotype III subtype calls for 16 genomes, all within serotype III. These were collapsed to III, the unit used in the literature, which is why the serotype counts sum to 83 while the raw output has 99 lines.

What the replicate comparison measures

S1 and S2 were sequenced together on one flow cell and basecalled in one run. They are not a partition of a single read set: three random partitions of the pooled reads give read N50 within 6 bp of each other, while S1 and S2 differ by 392 bp. The comparison therefore measures library preparation, sequencing and basecalling together. No barcode tag survives in the delivered FASTQ files, so the demultiplexing step itself could not be recovered from the data. The two assemblies differ at five positions. Three are substitutions at which both read sets carry both bases; because both libraries come from colony sweeps rather than single-colony picks, a mixture of two co-resident clones was a competing explanation. A mixture has one ratio, so every differentiating site must show the same minor-allele fraction. Pooled across both libraries the fractions are 29.4%, 46.6% and 55.8%, heterogeneous at chi-square = 59.25, 2 df, p = 1.4e-13, with the same rank order in each library separately. That excludes a fixed-ratio two-clone mixture and leaves context-dependent basecalling error, which is deterministic given the local sequence and recurs in both runs. Read-level phasing could not be used: the three positions lie 260 kb, 596 kb and 856 kb apart and no read spans a pair. Counts are in Supplementary Table S11.

What the comparison genomes are, and are not

The 83 comparison genomes are colonising isolates from rectovaginal swabs of near-term pregnant women at one hospital in Santo Domingo, sampled between September 2021 and March 2022. They are not invasive isolates and not a national sample. The sequenced isolate is a clinical isolate from a symptomatic vaginal infection in an adult patient, and is not among them. Prevalence figures describe that collection.

What the element boundaries rest on

Both elements are delimited from the outermost annotated mobility and conjugation genes flanking their cargo. Attachment sites, direct repeats and excision were not determined experimentally, so the sizes of approximately 34.4 kb and 35 kb are approximate. The tRNA-Lys adjacency fixes one boundary of element 2 more confidently than annotation alone.

When AMRFinderPlus cannot name an allele

AMRFinderPlus returns a family-level symbol when a hit does not meet its allele threshold, and such a symbol is not distinguishable from an allele in the two-column summary table. One genome, GCA_034336745.1 (ST2115/II), returned the bare symbol "lsa" twice, on separate contigs. The full output shows these are not partial or truncated genes: each covers 100% of the 492-residue reference at 90.0% identity to Lsa(C), so they are divergent Lsa-family ribosomal protection proteins that match no catalogued allele. They are the only determinants in the 83 genomes without an allele call. The per-genome AMRFinderPlus output is deposited in the code repository (amr83/), because the summary table alone cannot support this distinction.
